## Supplementary Information for "Cell-type-specific resolution epigenetics without the need for cell sorting or single-cell biology"

### 1 Supplementary Figures

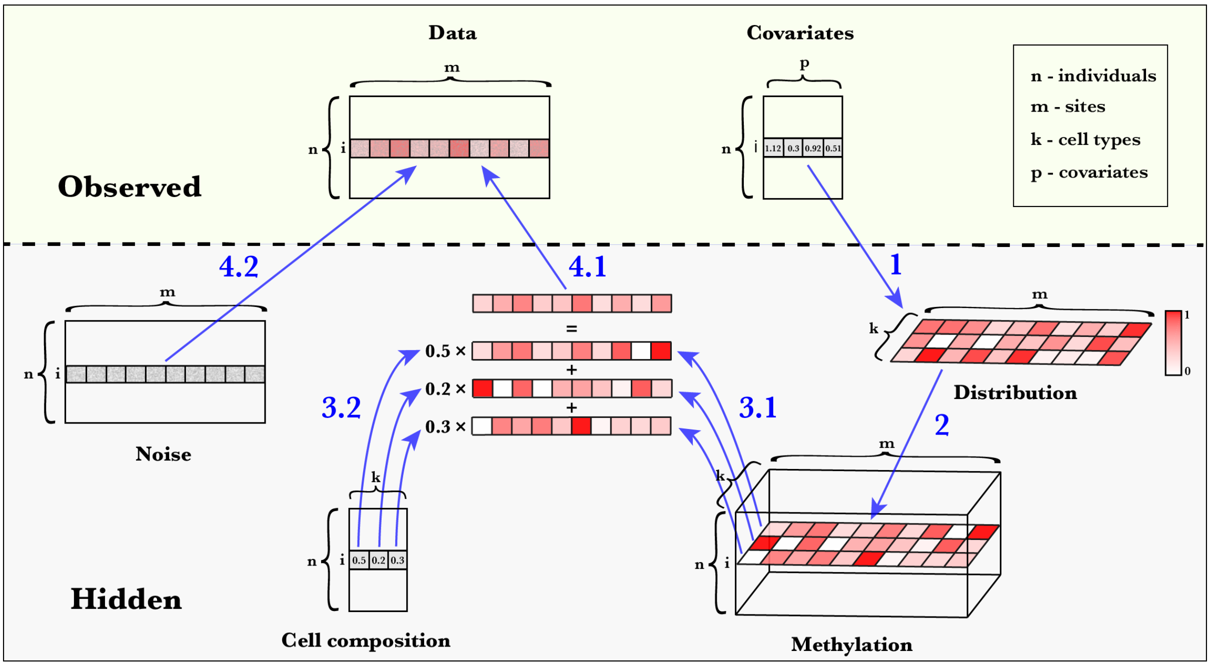

Figure 1: A summary of the TCA model for bulk DNA methylation data, presented as a four-steps generative model. Step 1: methylation altering covariates (e.g., age and sex) of a particular individual $i$ can affect the methylation distribution of individual $i$. Step 2: the cell-type-specific methylomes of individual $i$ are generated for each of the $k$ cell types in the studied tissue. Step 3: the cell-type-specific methylomes of individual $i$ (3.1) are combined according to the cell-type composition of the individual (3.2). Step 4: the true signal of the heterogeneous mixture (4.1) is distorted due to additional variation introduced by different sources of noise such as batch effects and other experiment-specific artificial variability (4.2); this results in the observed data. Methylation levels are represented by a gradient of red color

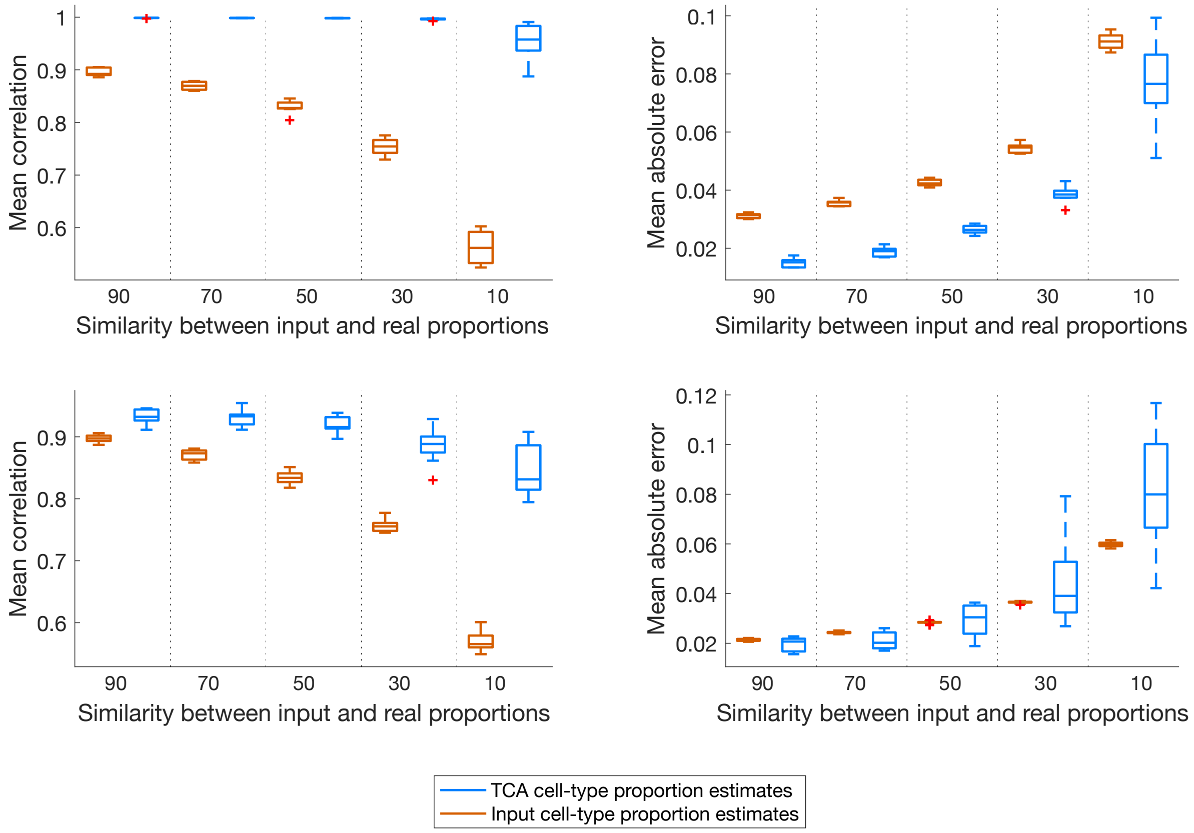

Figure 2: TCA improves noisy initial estimates of the cell-type proportions under different noise levels. For each noise level (induced by a level of similarity between the input and the real cell-type proportions; see Methods) of the input estimates, boxplots reflect the correlation and absolute error (averaged across cell-types) of the true cell-type proportions with the input estimates and with the TCA estimates. Results are presented under the assumption of three constituting cell types (k=3; top row) and under the assumption of six constituting cell types (k=6; bottom row), and each boxplot demosntrates the performance across 10 simulated datasets (the central mark on each box indicates the median, and the bottom and top edges indicate the 25th and 75th percentiles, respectively).

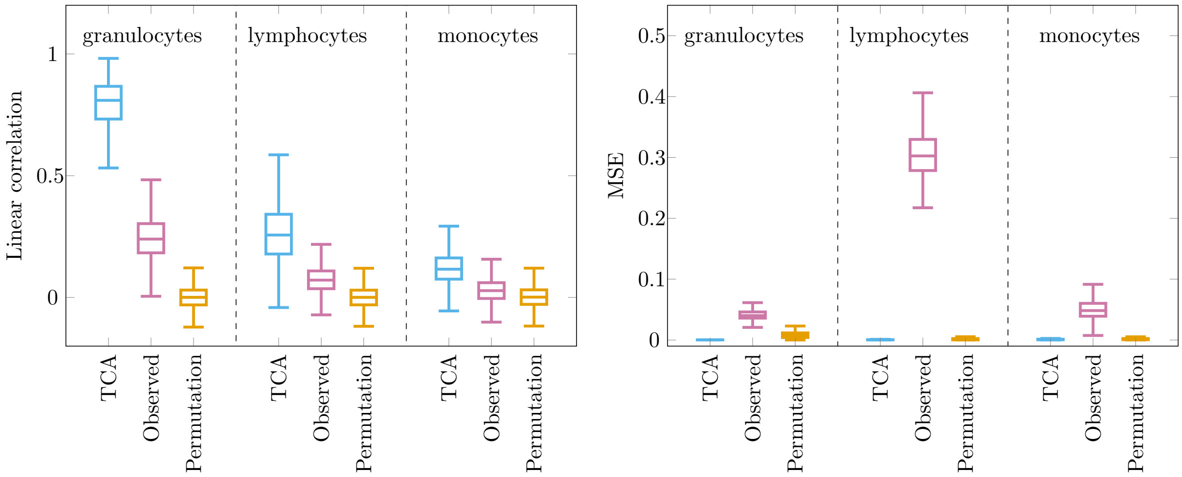

Figure 3: Reconstructing cell-type-specific methylation levels from simulated bulk whole-blood data with three constituting cell types ($k = 3$; 250 samples, 250 sites). Three approaches were evaluated in capturing the cell-type-specific levels of each site $j$ and cell type $h$ across all individuals $z_{hj} = (z_{hj}^1, ..., z_{hj}^n)$: TCA, TCA after permuting the observed data matrix ("Permutation") and directly using the observed bulk data ("Observed"; i.e. using the bulk as the estimate for the cell-type-specific levels of each cell type). For each of the evaluated approaches and for each of the simulated cell types (ordered by their mean abundance), presented are the distributions of the linear correlation between $z_{hj}$ and its estimate $\hat{z}_{hj}$ across all sites $j$ and across ten simulated data sets (left), and the distribution of the MSE between $z_{hj}$ and its estimate $\hat{z}_{hj}$ across all sites $j$ and across ten simulated data set (right). The central mark on each box indicates the median, and the bottom and top edges indicate the 25th and 75th percentiles, respectively.

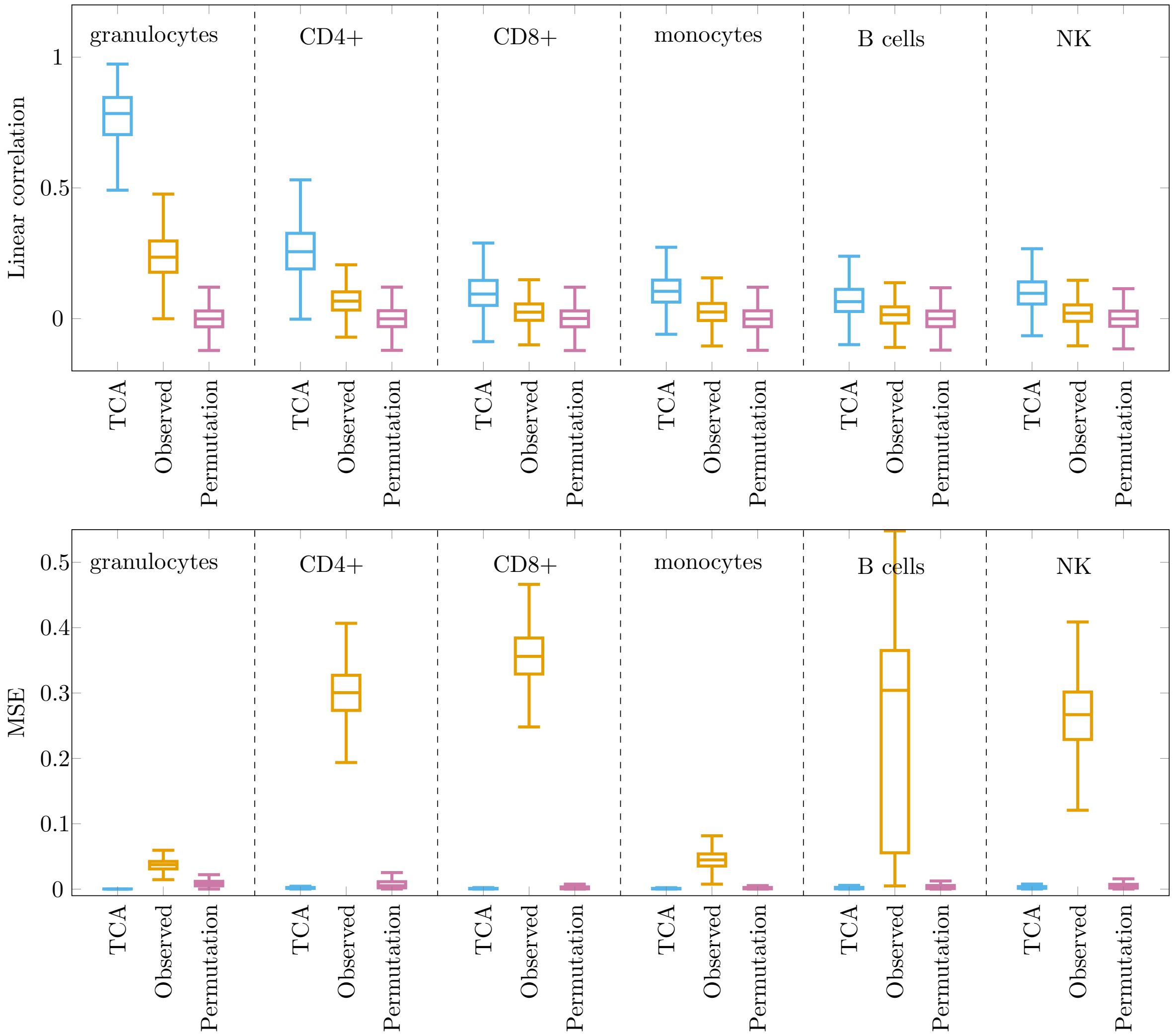

Figure 4: Reconstructing cell-type-specific methylation levels from simulated bulk whole-blood data with six constituting cell types ($k = 6$; 250 samples, 250 sites). Three approaches were evaluated in capturing the cell-type-specific levels of each site $j$ and cell type $h$ across all individuals $z_{hj} = (z^1_{hj}, ..., z^n_{hj})$: TCA, TCA after permuting the observed data matrix ("Permutation") and directly using the observed bulk data ("Observed"; i.e. using the bulk as the estimate for the cell-type-specific levels of each cell type). For each of the evaluated approaches and for each of the simulated cell types (ordered by their mean abundance), presented are the distributions of the linear correlation between $z_{hj}$ and its estimate $\hat{z}_{hj}$ across all sites $j$ and across ten simulated data sets (top), and the distribution of the MSE between $z_{hj}$ and its estimate $\hat{z}_{hj}$ across all sites $j$ and across ten simulated data set (bottom). The central mark on each box indicates the median, and the bottom and top edges indicate the 25th and 75th percentiles, respectively.

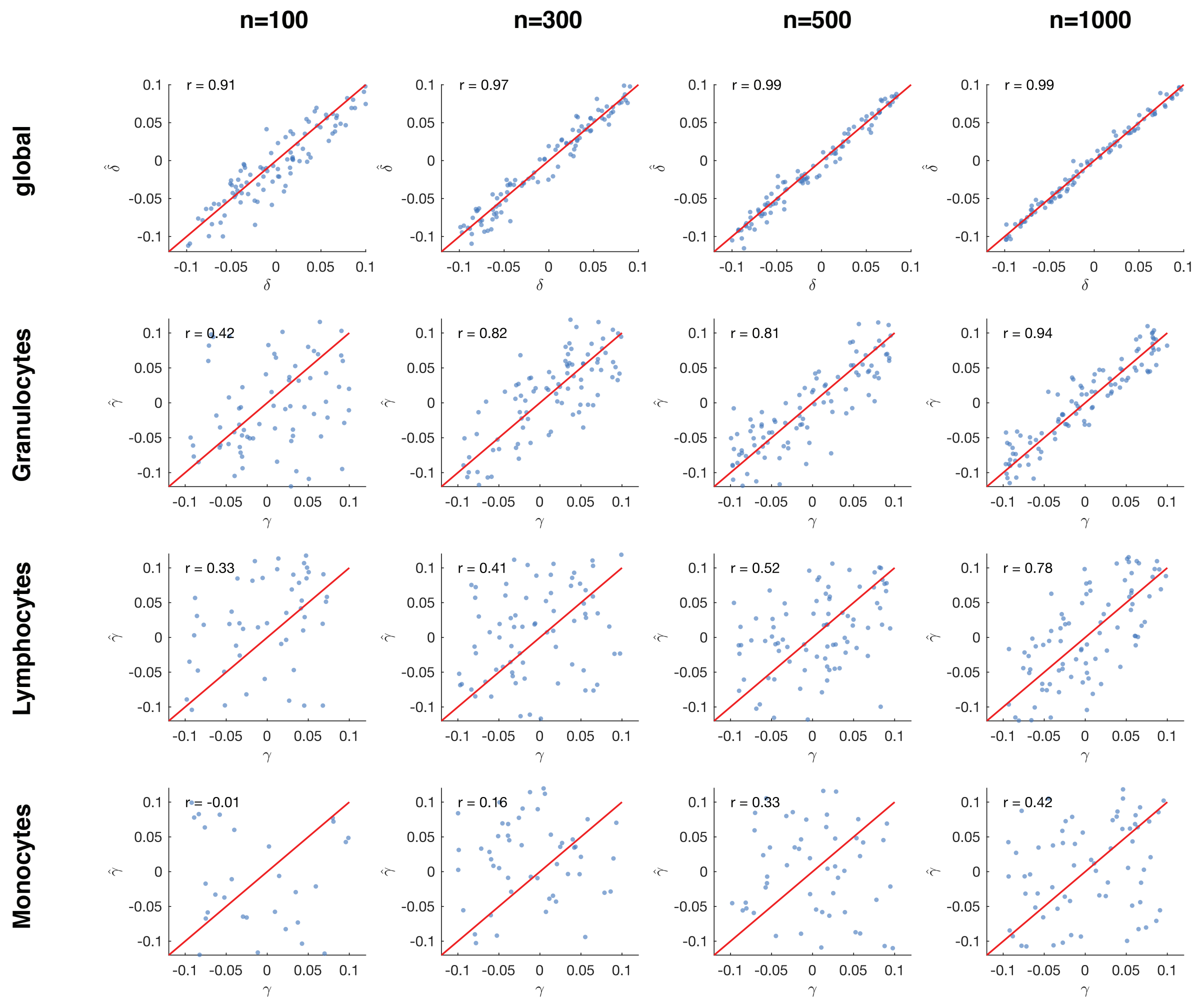

Figure 5: Estimating the effect sizes of covariates affecting methylation using TCA. Presented are true (X axes) and estimated (Y axes) effect sizes in simulated whole-blood methylation data with three constituting cell types ($k = 3$) and varying sample sizes (separated by different columns). Two scenarios were considered using a range of effect sizes: (1) estimating the effect of a covariate with global (i.e. non-cell-type-specific) effect on methylation (top row), and (2) estimating the effect of a covariate with cell-type-specific effect on methylation (rows 2-4). In the latter scenario, we considered a separate experiment for each of the three cell types, such that in each experiment the covariate affected the particular cell type under test. Throughout these experiments, covariates were generated from a normal distribution, and both global ($\delta$) and cell-type-specific ($\gamma$) effect sizes were sampled from a uniform distribution.

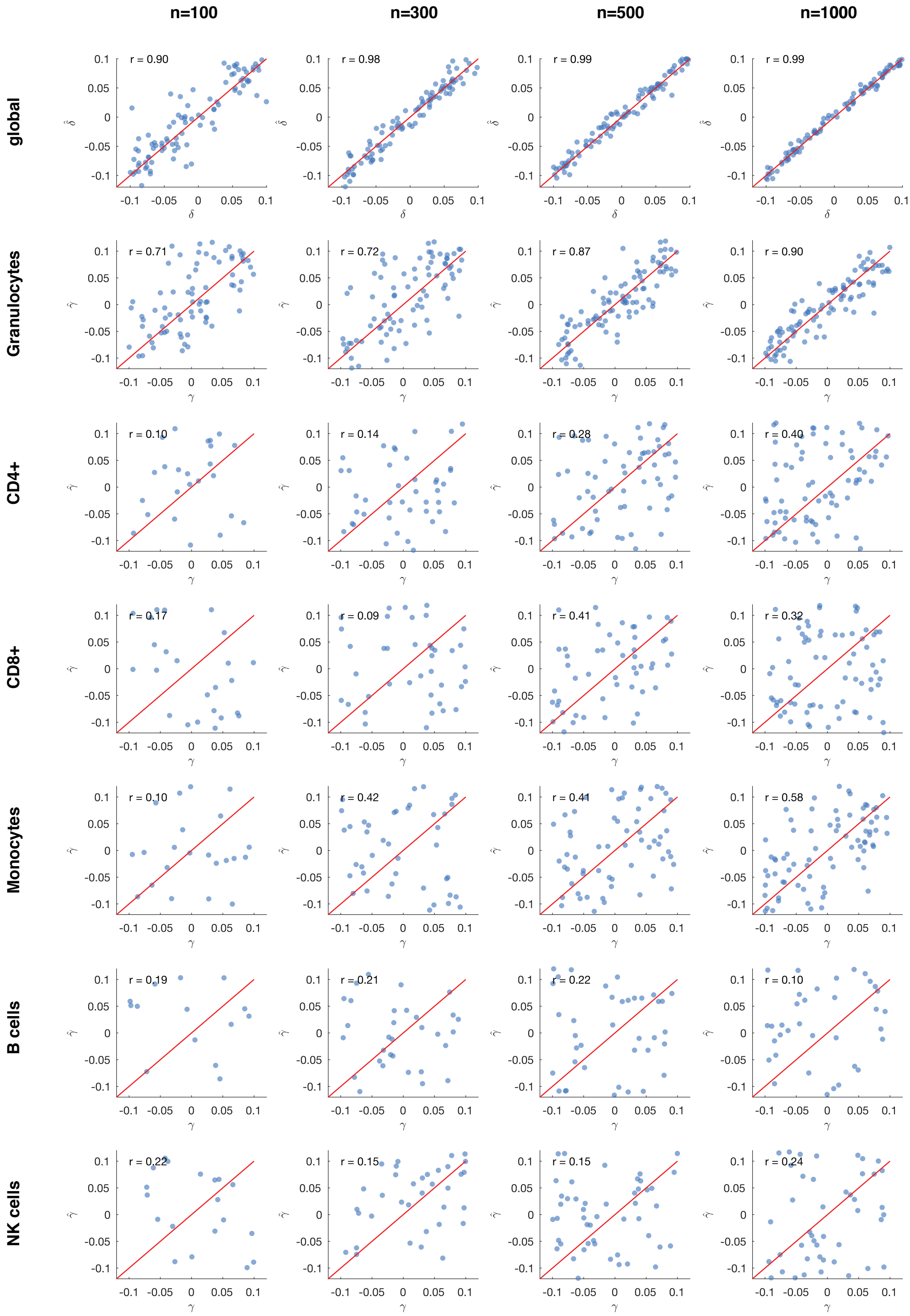

n=100
n=300
n=500
n=1000
global
r = 0.90
r = 0.98
r = 0.99
r = 0.99
Granulocytes
r = 0.71
r = 0.72
r = 0.87
r = 0.90
CD4+
r = 0.10
r = 0.14
r = 0.28
r = 0.40
CD8+
r = 0.17
r = 0.09
r = 0.41
r = 0.32
Monocytes
r = 0.10
r = 0.42
r = 0.41
r = 0.58
B cells
r = 0.19
r = 0.21
r = 0.22
r = 0.10
NK cells
r = 0.22
r = 0.15
r = 0.15
r = 0.24

Figure 6: Estimating the effect sizes of covariates affecting methylation using TCA. Presented are true (X axes) and estimated (Y axes) effect sizes in simulated whole-blood methylation data with six constituting cell types ($k = 6$) and varying sample sizes (separated by different columns). Two scenarios were considered using a range of effect sizes: (1) estimating the effect of a covariate with global (i.e. non-cell-type-specific) effect on methylation (top row), and (2) estimating the effect of a covariate with cell-type-specific effect on methylation (rows 2-7). In the latter scenario, we considered a separate experiment for each of the three cell types, such that in each experiment the covariate affected the particular cell type under test. Throughout these experiments, covariates were generated from a normal distribution, and both global ($\delta$) and cell-type-specific ($\gamma$) effect sizes were sampled from a uniform distribution.

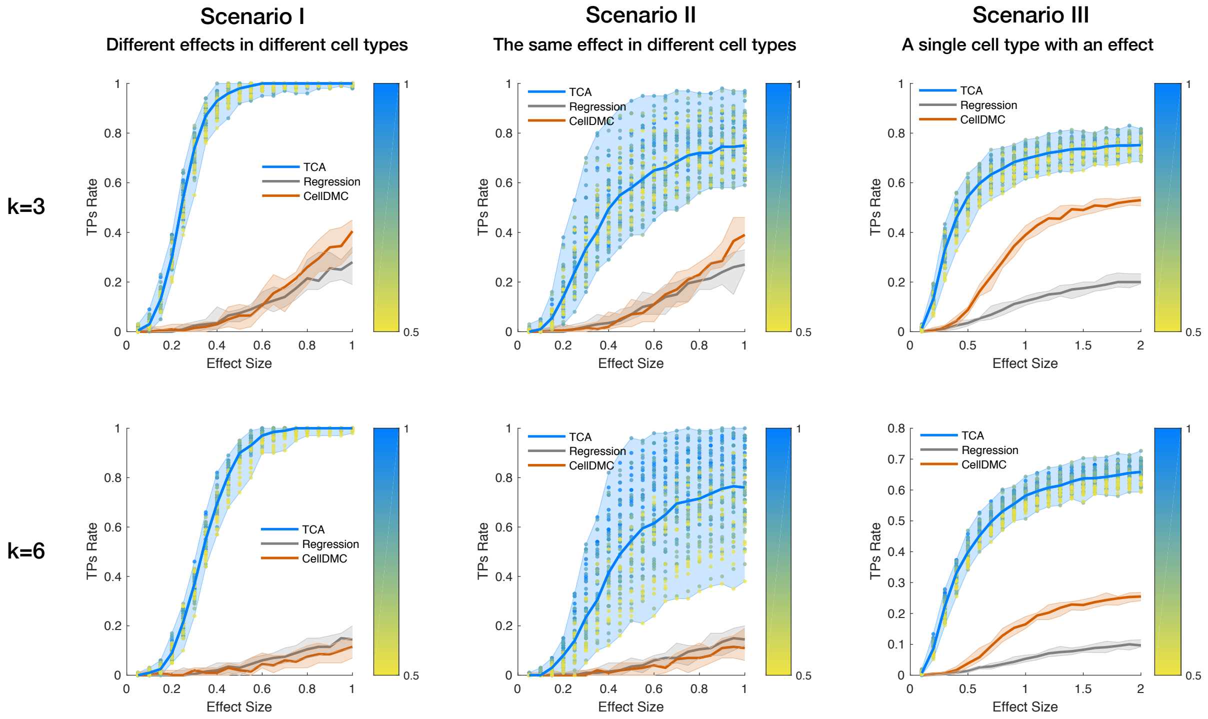

Figure 7: An evaluation of power for detecting cell-type-specific associations with DNA methylation while including cell-type-specific affecting covariates and using a nonparametric distribution of the cell-type proportions. Performance was evaluated using three approaches: TCA, a standard linear regression with the observed bulk data, and CellDMC with the true cell-type proportions as an input. The numbers of true positives (TPs) were measured under three scenarios using a range of effect sizes: different effect sizes for different cell types (Scenario I), the same effect size for all cell types (Scenario II), and a single effect size for a single cell type (Scenario III); each of the scenarios was evaluated under the assumption of three constituting cell types (k=3; top row) and six constituting cell types (k=6; bottom row). Lines represent the median performance across 10 simulations and the colored areas reflect the results range across the multiple executions. The colored dots reflect the results of TCA under different initializations of the cell-type proportion estimates (i.e. different levels of noise injected into TCA), where the color gradients represent the mean absolute correlation of the initial estimates with the true values (across all cell types).

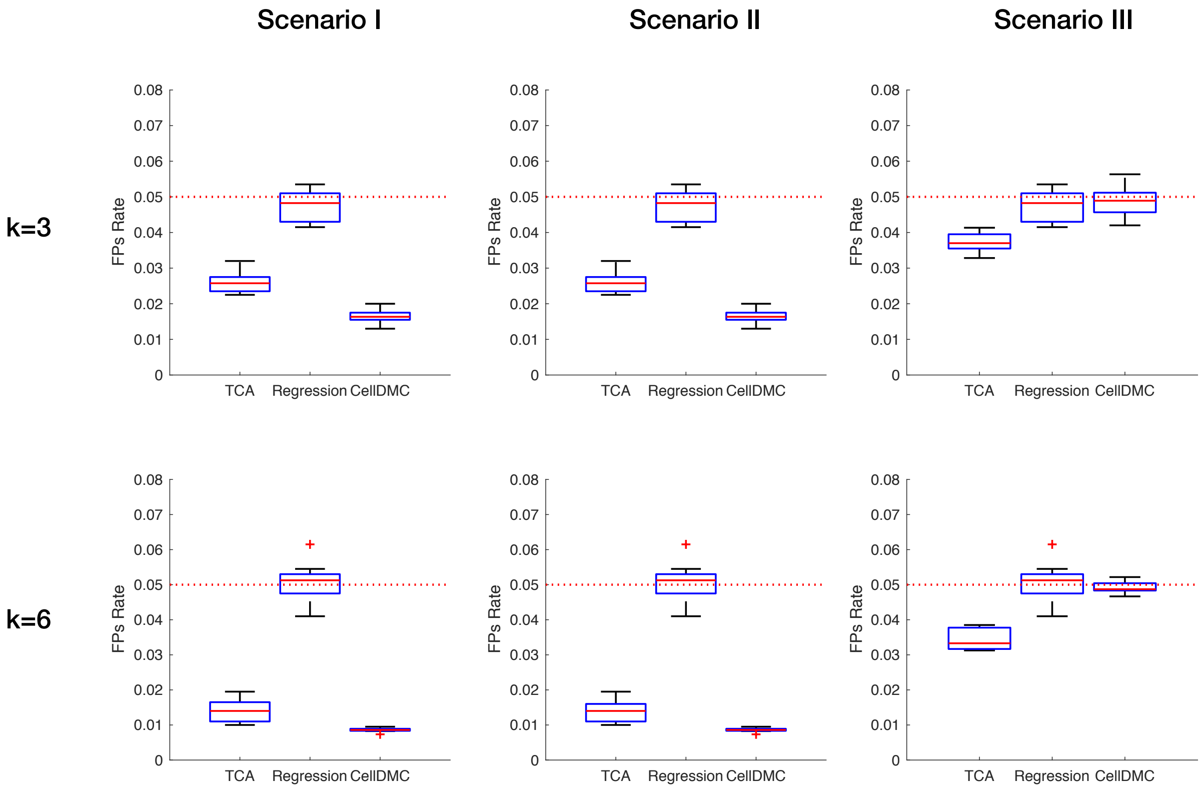

Figure 8: An evaluation of false positives rates in association testing with DNA methylation. Performance was evaluated using three approaches: TCA, a standard linear regression with the observed bulk data, and CellDMC with the true cell-type proportions as an input. The proportions of false positives (FPs) were measured under three scenarios using a range of effect sizes: different effect sizes for different cell types (Scenario I), the same effect size for all cell types (Scenario II), and only a single effect size for a single cell type (Scenario III); each of the scenarios was evaluated under the assumption of three constituting cell types (k=3) and six constituting cell types (k=6). Boxplots reflect results across 10 simulations. The central mark on each box indicates the median, and the bottom and top edges indicate the 25th and 75th percentiles, respectively.

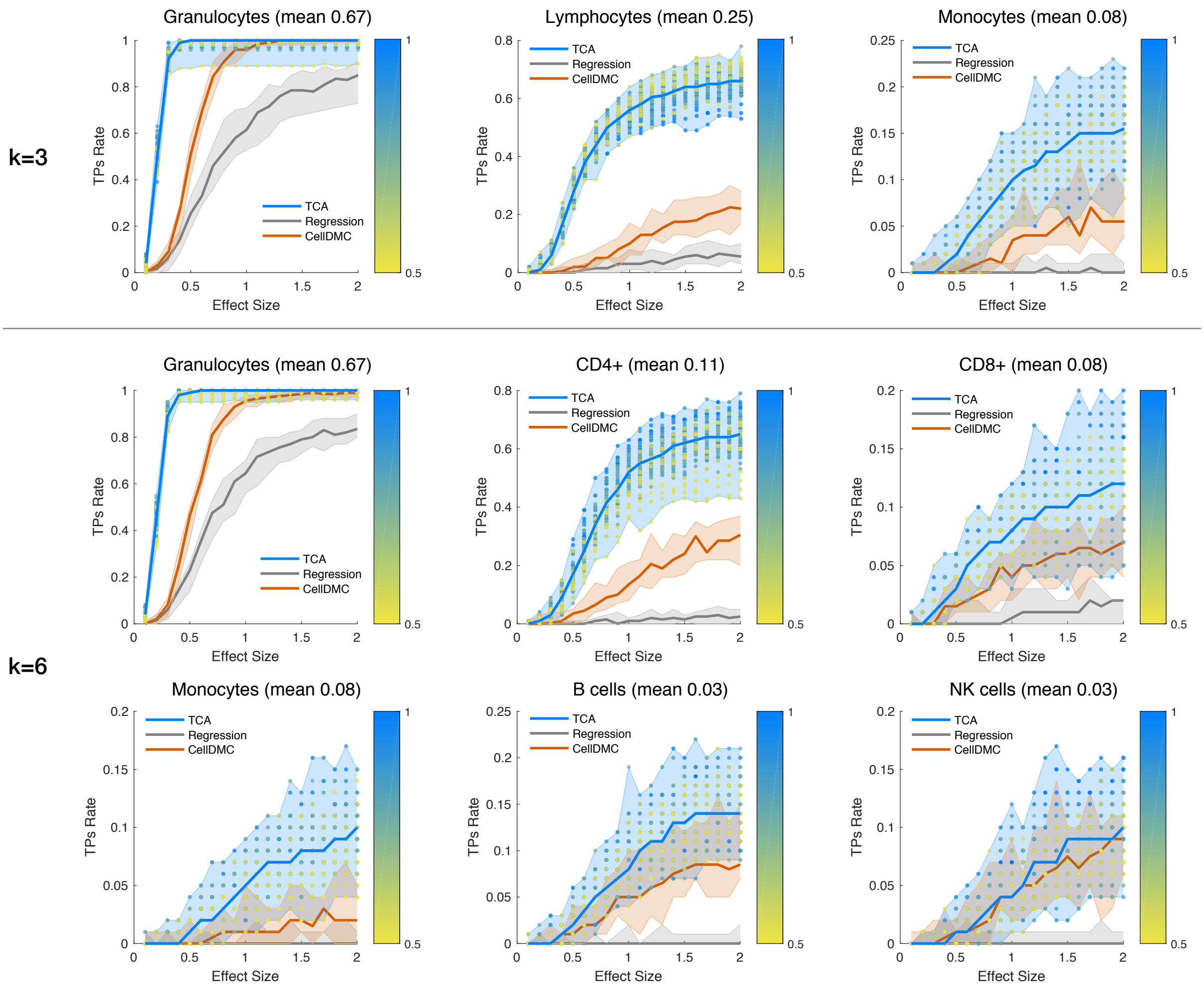

Figure 9: An evaluation of power for detecting cell-type-specific associations with DNA methylation, stratified by cell types (with the mean abundance of each cell type noted). Performance was evaluated using three approaches: TCA, a standard linear regression with the observed bulk data, and CellDMC with the true cell-type proportions as an input. The numbers of true positives were measured under a scenario where only a single effect size for a single cell type exists, both in the case of three constituting cell types (k=3) and six constituting cell types (k=6). The colored areas reflect the results range across 10 simulations, and the colored dots reflect the results of TCA under different initializations of the cell-type composition estimates (i.e. different levels of noise injected into TCA), where the color gradients represent the mean absolute correlation of the initial estimates with the true values (across all cell types).

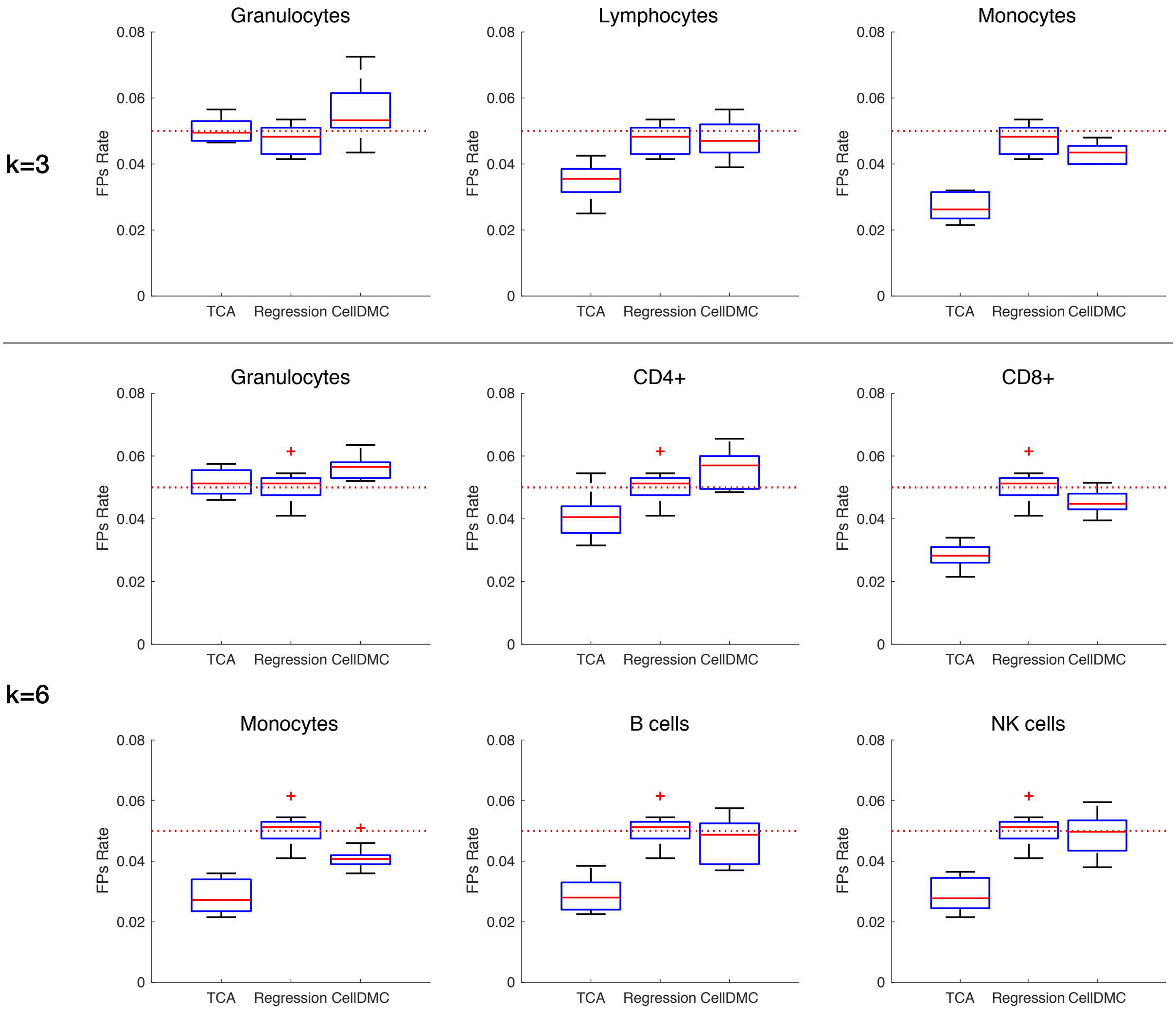

Figure 10: An evaluation of false positives rates in association testing with DNA methylation, stratified by cell types. Performance was evaluated using three approaches: TCA, a standard linear regression with the observed bulk data, and CellDMC with the true cell-type proportions as an input. The proportions of false positives (FPs) were measured under a scenario where only a single effect size for a single cell type exists, both in the case of three constituting cell types (k=3) and six constituting cell types (k=6). Boxplots reflect results across 10 simulations. The central mark on each box indicates the median, and the bottom and top edges indicate the 25th and 75th percentiles, respectively.

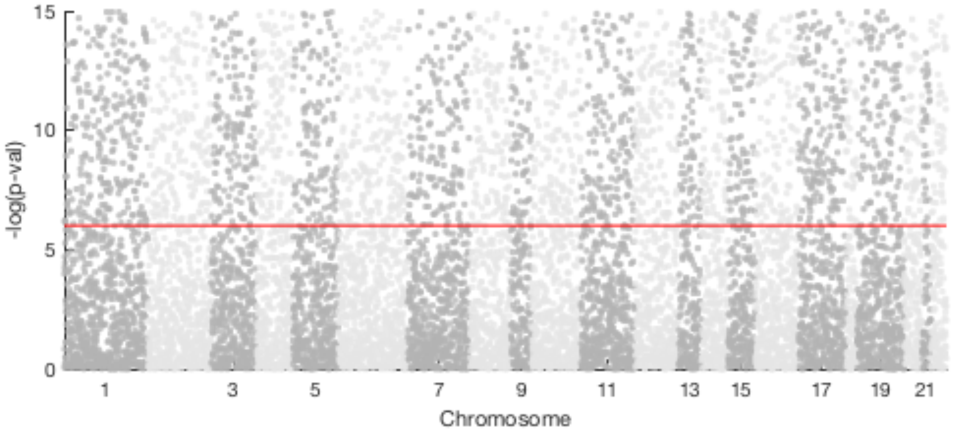

Figure 11: Results of the association analysis with level of immune activity using a standard regression model. Presented is a Manhattan plot of the -log10 P-values for the association tests (results subsampled and truncated for visualization), where the horizontal red line represents the experiment-wide significance threshold.

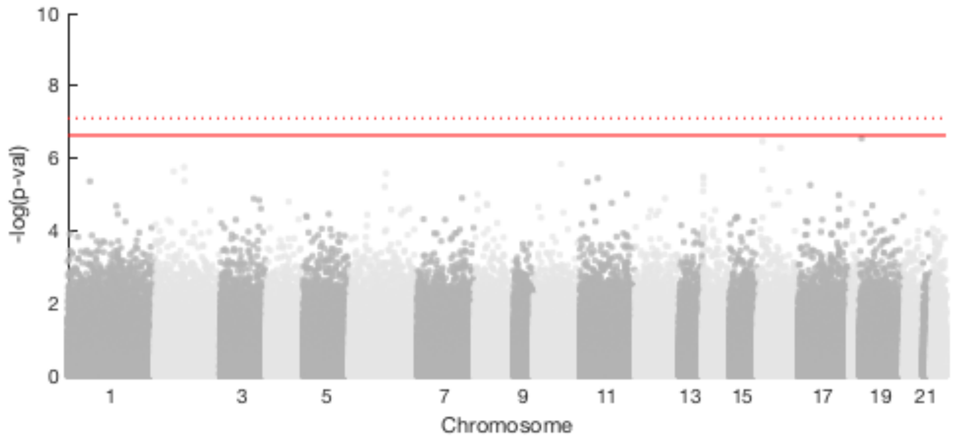

Figure 12: Results of the cell-type-specific association analysis with RA using a standard regression model in the Rhead et al. sorted methylation data. Presented is a Manhattan plot of the -log10 P-values for the association tests in CD4+, CD14+, and CD+19 cells. The solid horizontal red line represents the experiment-wide significance threshold, and the dotted horizontal red line represents the significance threshold adjusted for three experiments corresponding to the three cell types.

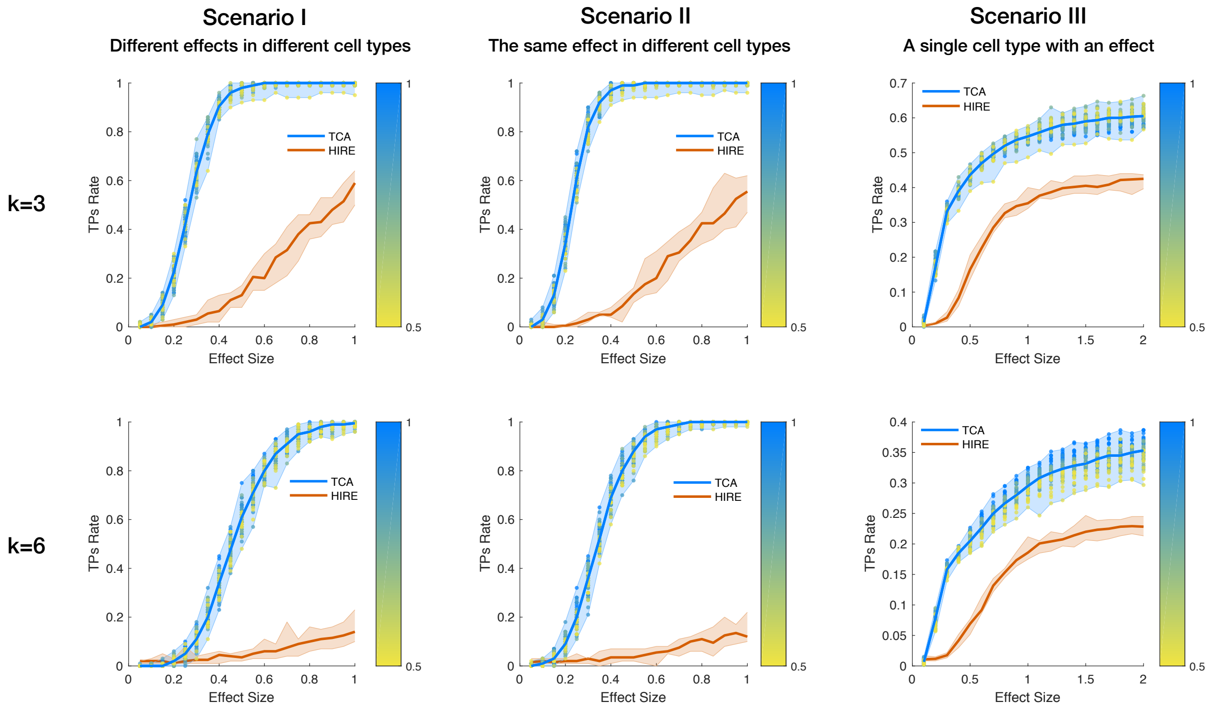

Figure 13: An evaluation of power for detecting cell-type-specific associations with DNA methylation. Performance was evaluated using two approaches: TCA and HIRE by Luo et al. with the true cell-type proportions as an input. The numbers of true positives (TPs) were measured under three scenarios using a range of effect sizes: different effect sizes for different cell types (Scenario I), the same effect size for all cell types (Scenario II), and a single effect size for a single cell type (Scenario III); each of the scenarios was evaluated under the assumption of three constituting cell types (k=3; top row) and six constituting cell types (k=6; bottom row). Lines represent the median performance across 10 simulations and the colored areas reflect the results range across the multiple executions. The colored dots reflect the results of TCA under different initializations of the cell-type proportion estimates (i.e. different levels of noise injected into TCA), where the color gradients represent the mean absolute correlation of the initial estimates with the true values (across all cell types).

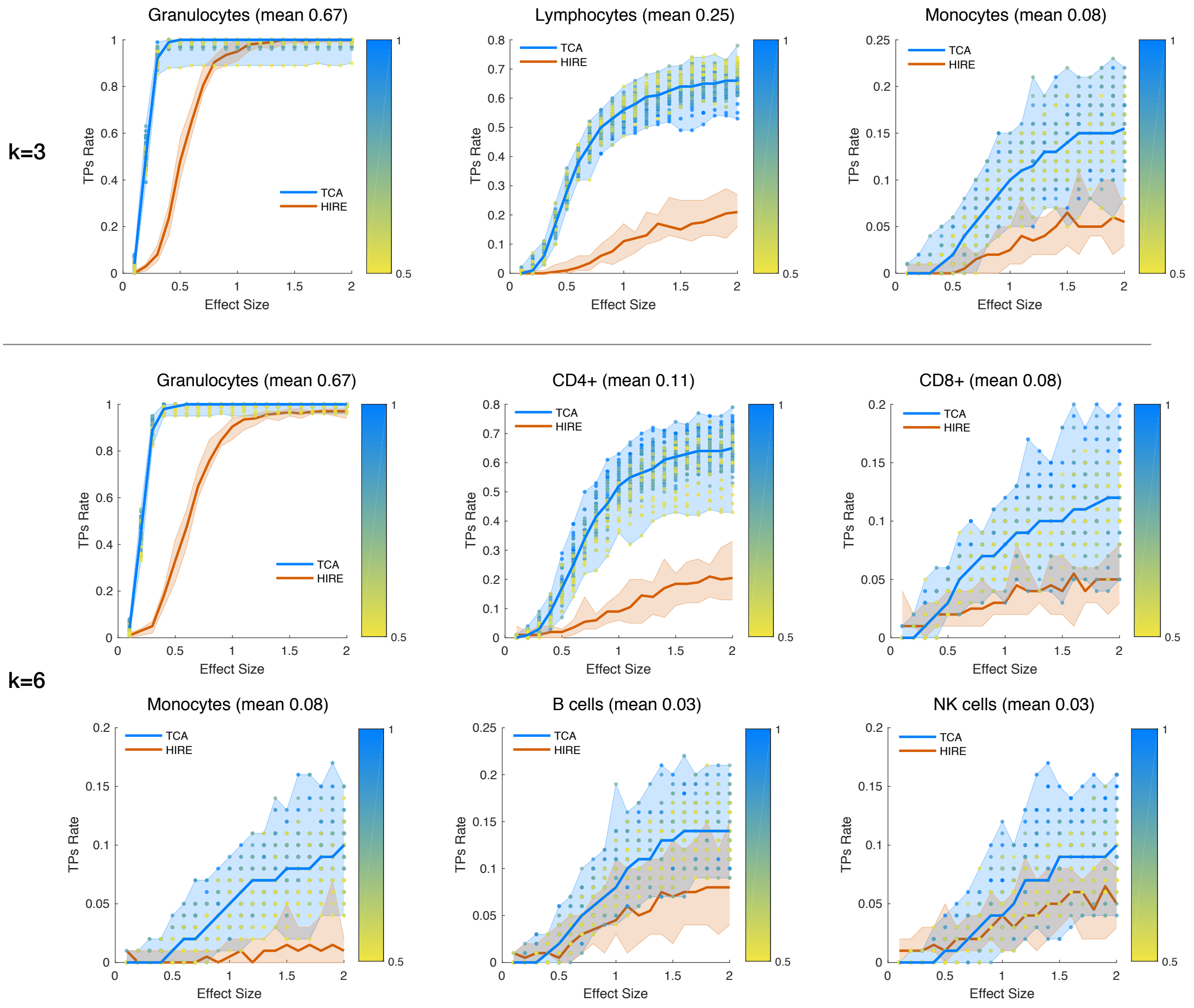

Figure 14: An evaluation of power for detecting cell-type-specific associations with DNA methylation, stratified by cell types (with the mean abundance of each cell type noted). Performance was evaluated using two approaches: TCA and HIRE by Luo et al. with the true cell-type proportions as an input. The numbers of true positives were measured under a scenario where only a single effect size for a single cell type exists, both in the case of three constituting cell types (k=3) and six constituting cell types (k=6). The colored areas reflect the results range across 10 simulations, and the colored dots reflect the results of TCA under different initializations of the cell-type composition estimates (i.e. different levels of noise injected into TCA), where the color gradients represent the mean absolute correlation of the initial estimates with the true values (across all cell types).

### 2 Supplementary Methods

#### 2.1 Technical background

In a classical matrix factorization problem, a matrix $X \in \mathbb{R}^{n \times m}$ is given and we look for two matrices $W \in \mathbb{R}^{k \times n}, Z \in \mathbb{R}^{k \times m}$ such that $X \approx W^T Z$. The matrix $W$ is commonly referred to as the weights matrix, and $Z$ is commonly referred to as the features matrix. The dimension parameter of the factorization (decomposition) $k$ is typically known a-priori or assumed to be known. This factorization problem is typically formulated as an optimization problem, in which we aim at finding the values of $W, Z$ that minimize some loss function with respect to $X$ (e.g., the Frobenius norm). Put differently, we look for a decomposition $X = W^T Z + E$, such that the error matrix $E \in \mathbb{R}^{n \times m}$ is minimized in some sense.

In a typical setup, $x_{ij}$ denotes the $i$-th observation of the $j$-th feature of $X$ for some type of measurement, and its factorization is considered to be $w_i$ (the $i$-th column of $W$) and $z_j$ (the $j$-th column of $Z$). Therefore, $x_{ij}$ can be interpreted as a linear combination of $k$ different sources of some feature $j$, weighted according to the $k$ observation-specific weights in $w_i$. Introducing constraints on $W$ or $Z$ (or both) is often applied in many real-world problems; for example, Principal Component Analysis (PCA) and Factor Analysis are two widely-used methods that constrain the rows of $W, Z$ to be orthogonal, and can further be extended to penalize their norms in order to induce desired properties such and sparsity (e.g., [1]).

Various extensions of matrix factorization that assume explicit generative models for $X$ exist. Notably, probabilistic approaches to matrix factorization (PMF) are used to infer knowledge about assumed distributions over $W, Z$, an approach which can be further extended by assuming priors on those distributions (e.g., [2,3]). Such PMF formulations typically infer a point estimate of the likelihood or posterior of $W, Z$ or try to infer their full posterior distribution.

Both the matrix factorization problem and the PMF problem essentially assume that the features matrix $Z$ is shared across observations (albeit the latter assumes that $Z$ is coming from a distribution). Here, we propose a model where $x_{ij}$ is a linear combination of $w_i, z_j^i$, where

$z^i_j = (z^i_{1j}, ..., z^i_{kj})^T$ is coming from a vector of random variables $Z^i_j$, representing $k$ sources of feature $j$ that are unique to observation $i$. This model comes with a statistical challenge, as the number of parameters exceeds the number of observations. To enable inference, we require $Z^i_j$ to be random but to share parameters across observations.

We present a generalization of the standard matrix factorization problem, which considers a generative model with the assumption that each observation may have a unique features matrix. Unlike previous formulations of matrix factorization, where we assume that the mean is factorized, our model assumes that both the mean and the variance are factorized. As a result, as we show, knowledge of $W$ and the distribution of $\{Z^i_j\}$ (which can be estimated) allows us to extract information about the $\{z^i_{hj}\}$ values. In our context of DNA methylation, these values represent the underlying methylation levels for each sample $i$ in each cell type $h$ and at each site $j$. Below, we first describe the model and a procedure for estimating the $\{z^i_{hj}\}$ values, which together compose a three-dimensional tensor. Then, we consider the problem of detecting cell-type-specific associations in epigenetic studies, and we derive a direct statistical test based on our proposed model.

#### 2.2 Tensor Composition Analysis (TCA): Intuition

Unlike existing factorization models for DNA methylation [4–8], which explicitly (or implicitly) assume that all individuals have the same cell-type-specific methylation profiles (methylomes), in our model each individual has unique cell-type-specific methylomes. Since cell-type-specific genomic features are expected to demonstrate inherent biological variability across individuals, we argue that this model is a more realistic one compared with the alternative factorization models.

While the presumption of non-systematic individual- and cell-type-specific variability justifies this model, some of the variability in a given cell type may be systematic. We therefore consider the potential systematic effects of covariates (e.g., age and gender) on the cell-type-specific variability. In addition, we consider the potential effect of covariates that do not have cell-type-specific effects but rather may have direct effects on the observed mixtures (e.g., batch effects).

Below, we present the TCA model in a general way, which takes potential covariates into account. We derive the conditional probability of the unobserved individual- and cell-type-specific methylation levels given the observed methylation mixtures, which can then be used for estimating cell-type-specific levels for each individual, and we demonstrate mathematically and empirically the reason why TCA works. Since TCA requires knowledge of the parameters of the model (which are typically not available in practice) we further describe a maximum-likelihood (ML) based procedure for inferring the parameters of the model based on the observed methylation levels. We further derive a one-step statistical test for association testing between a phenotype of interest and cell-type-specific methylation levels by deriving the conditional distribution of the phenotype given the observed methylation mixtures. Finally, we provide insights into the reason why applying a standard regression approach to bulk data is severely biased, and show that TCA provides an unbiased solution.

#### 2.3 The TCA model

Let $Z_{hj}^i$ be the methylation level of individual $i \in \{1, ...n\}$ in cell type $h \in \{1, ...k\}$ at methylation site $j \in \{1, ...m\}$, and let $C^{(1)} \in \mathbb{R}^{p_1 \times n}$ be a matrix of $p_1$ covariates that may potentially affect methylation levels in a cell-type-specific manner. We assume:

$$Z_{hj}^i = \mu_{hj} + (c_i^{(1)})^T \gamma_h^j + \epsilon_{hj}^i \tag{1}$$

$$\epsilon_{hj}^i \sim N(0, \sigma_{hj}^2) \tag{2}$$

where $c_i^{(1)}$ is the $i$-th column of $C^{(1)}$ (corresponding to the $p_1$ covariates of the $i$-th individual), $\gamma_h^j$ is a $p_1$-length vector of corresponding effects sizes for the $p_1$ covariates in the $h$-th cell type at site $j$, and $e_{hj}^i$ is an i.i.d. component of variation.

We assume that observed methylation levels are convolved signals coming from $k$ different cell-types. We denote $W \in \mathbb{R}^{k \times n}$ as a matrix of cell-type proportions of $k$ cell types for each of the $n$ individuals, and $C^{(2)} \in \mathbb{R}^{p_2 \times n}$ as a matrix of $p_2$ global covariates potentially affecting the observed methylation levels. Our model for $X_{ij}$, the observed methylation level of the $i$-th individual in cell type $j$, is as follows:

$$\begin{aligned} X_{ij} &= (c_i^{(2)})^T \delta_j + \sum_{h=1}^{k} w_{hi} Z_{hj}^i + \epsilon_{ij} & (3)\\ & \epsilon_{ij} \sim N(0, \tau^2) & (4)\\ \text{s.t.} \quad & \forall i : \sum_{h=1}^{k} w_{hi} = 1 & (5)\\ & \forall h, i : w_{hi} \geq 0 & (6) \end{aligned}$$

where $c_i^{(2)}$ is the $i$-th column of $C^{(2)}$ (corresponding to the $p_2$ covariates of the $i$-th individual), $\delta_j$ is a $p_2$-length vector of corresponding effects sizes of the $p_2$ covariates for the $j$-th site, and $e_{ij}$ is a component of i.i.d. variation that models measurement noise.

##### 2.4 Deriving the TCA estimator

Let $\Theta_j = (\mu_j, \sigma_j, w_i, \tau, \Gamma_j, \delta_j)$ be the set of the model's parameters for a particular site $j$, where $\Gamma_j$ is a $p_1 \times k$ matrix with the vectors $\gamma_1^j, ..., \gamma_k^j$. Given the observed values, we are interested in the conditional distribution $Z_j^i | X_{ij} = x_{ij}$. Following the assumptions in (1) to (4), the conditional probability satisfies:

$$\begin{aligned} &Pr(Z_j^i = z_j^i | X_{ij} = x_{ij}, c_i^{(1)}, c_i^{(2)}, \Theta_j) \propto Pr(Z_j^i = z_j^i | \mu_j, \sigma_j, c_i^{(1)}, \Gamma_j) Pr(X_{ij} = x_{ij} | Z_j^i = z_j^i, w_i, \tau, c_i^{(2)}, \delta_j)\\ &\quad \propto exp\left(-\frac{1}{2}\left(z_j^i - \mu_j - \Gamma_j^T c_i^{(1)}\right)^T \Sigma_j^{-1} \left(z_j^i - \mu_j - \Gamma_j^T c_i^{(1)}\right)\right)\\ &\quad exp\left(-\frac{1}{2\tau^2}\left(x_{ij} - (z_j^i)^T w_i - (c_i^{(2)})^T \delta_j\right)^2\right)\\ &\quad \propto exp\left(-\frac{1}{2}\left((z_j^i)^T \Sigma_j^{-1} z_j^i - 2(z_j^i)^T \Sigma_j^{-1}\left(\mu_j + \Gamma_j^T c_i^{(1)}\right)\right)\right)\\ &\quad exp\left(-\frac{1}{2\tau^2}\left((z_j^i)^T w_i w_i^T z_j^i - 2(z_j^i)^T w_i\left(x_{ij} + (c_i^{(2)})^T \delta_j\right)\right)\right)\\ &\quad \propto exp\left(-\frac{1}{2}\left((z_j^i)^T\left(\Sigma_j^{-1} + \frac{w_i w_i^T}{\tau^2}\right) z_j^i\right)\right)\\ &\quad exp\left(-\frac{1}{2}\left(-2(z_j^i)^T\left(\Sigma_j^{-1}\left(\mu_j + \Gamma_j^T c_i^{(1)}\right) + w_i\left(\frac{x_{ij} + (c_i^{(2)})^T \delta_j}{\tau^2}\right)\right)\right)\right)\\ &\quad \propto exp\left(-\frac{1}{2}(z_j^i - a_{ij})^T S_{ij}^{-1}(z_j^i - a_{ij})\right) \end{aligned} \quad (7)$$

where

$$\Sigma_j = diag(\sigma_{1j}^2, ..., \sigma_{kj}^2) \tag{8}$$

$$S_{ij} = \left(\Sigma_j^{-1} + \frac{w_i w_i^T}{\tau^2}\right)^{-1} \tag{9}$$

$$a_{ij} = S_{ij}\left(\Sigma_j^{-1}\left(\mu_j + \Gamma_j^T c_i^{(1)}\right) + w_i\left(\frac{x_{ij} + (c_i^{(2)})^T \delta_j}{\tau^2}\right)\right) \tag{10}$$

The probability in (7) is maximized when $z_j^i$ is the mode of the conditional distribution (which is the mean in this case). We therefore set the TCA estimator of $z_j^i$ to be:

$$\hat{z}_j^i = a_{ij} = \left(\frac{w_i w_i^T}{\tau^2} + \Sigma_j^{-1}\right)^{-1}\left(\Sigma_j^{-1}\left(\mu_j + \Gamma_j^T c_i^{(1)}\right) + w_i\left(\frac{x_{ij} + (c_i^{(2)})^T \delta_j}{\tau^2}\right)\right) \tag{11}$$

#### 2.5 Extracting underlying signals from convolved signals using TCA

In order to see why TCA can learn non-trivial information about the $\{z_{hj}^i\}$ values, note that [9]

$$Z_{hj}^i | X_{ij} \sim N\left(\tilde{\mu}_1 + \frac{COV(Z_{hj}^i, X_{ij})}{\tilde{\sigma}_2^2}(x_{ij} - \tilde{\mu}_2), \tilde{\sigma}_1^2 - \frac{COV(Z_{hj}^i, X_{ij})^2}{\tilde{\sigma}_2^2}\right) \tag{12}$$

where

$$\tilde{\mu}_1 = E(Z_{hj}^i), \tilde{\sigma}_1^2 = V(Z_{hj}^i) \tag{13}$$

$$\tilde{\mu}_2 = E(X_{ij}), \tilde{\sigma}_2^2 = V(X_{ij}) \tag{14}$$

Consider a simplified case where $\tau = 0$ and $\mu_{hj} = 0, \sigma_{hj} = 1$ for each $h$ and some particular $j$. Assuming no covariates for simplicity, given the model of $Z_{hj}^i$ and the model of $X_{ij}$ in (1) to (4), we know that

$$\begin{aligned} COV(Z^i_{hj}, X_{ij}) &= E(Z^i_{hj} X_{ij}) - E(Z^i_{hj})E(X_{ij}) \\ &= E\left(Z^i_{hj} \sum_{l=1}^{k} w_{li} Z^i_{lj}\right) \\ &= E\left(w_{hi}(Z^i_{hj})^2 + E(Z^i_{hj}) \sum_{l \neq h} w_{li} E(Z^i_{lj})\right) \\ &= E\left(w_{hi}\left(V(Z^i_{hj}) + E(Z^i_{hj})^2\right)\right) \\ &= w_{hi} \end{aligned} \tag{15}$$

Therefore, based on (12), in this case we get that

$$Z^i_{hj} | X_{ij} = x_{ij} \sim N\left(\frac{w_{hi} x_{ij}}{\sum_{l=1}^{k} w_{li}^2}, 1 - \frac{w_{hi}^2}{\sum_{l=1}^{k} w_{li}^2}\right) \tag{16}$$

This means that given the observed value $x_{ij}$, the conditional distribution of $Z^i_{hj}$ has a lower variance compared with that of the marginal distribution of $Z^i_{hj}$ ($\sigma^2_{hj} = 1$), thus reducing the uncertainty and allowing us to provide a non-trivial estimate for the $\{z^i_{hj}\}$ values. This result is not specific for methylation but rather more general. In order to empirically verify this result and get an initial intuition as for the potential performance of TCA, we considered the following simplified general simulation.

We sampled three-dimensional source- and observation-specific values according to the model in (1)-(2) for every feature $j$, observation $i$ and source $h$ (i.e. for each of the $\{z^i_{hj}\}$ values) using $n = 250, m = 250, k = 3$ for the number of observations, features and sources, respectively. In this experiment, we sampled all the source- and observation-specific values, as well as the weights matrix ($W$), from a standard normal distribution. Eventually, we generated a matrix of observed mixtures ($X$) according to the model in (3)-(4) using the source- and observation-specific values, the weights matrix and an additional component of i.i.d. variation ($\tau = 0.01$). For performance evaluation, for each estimated vector $\hat{z}_{hj} = (\hat{z}^1_{hj}, ..., \hat{z}^n_{hj})^T$, we considered its linear correlation and mean squared error (MSE) with the true values in $z_{hj}$.

For simplicity, we assumed that all the parameters of the model are known, and applied TCA for estimating the $\{z^i_{hj}\}$ values. In order to form a baseline for comparison and to empirically verify that TCA can extract non-trivial information about the $\{z^i_{hj}\}$ values, we also applied TCA after

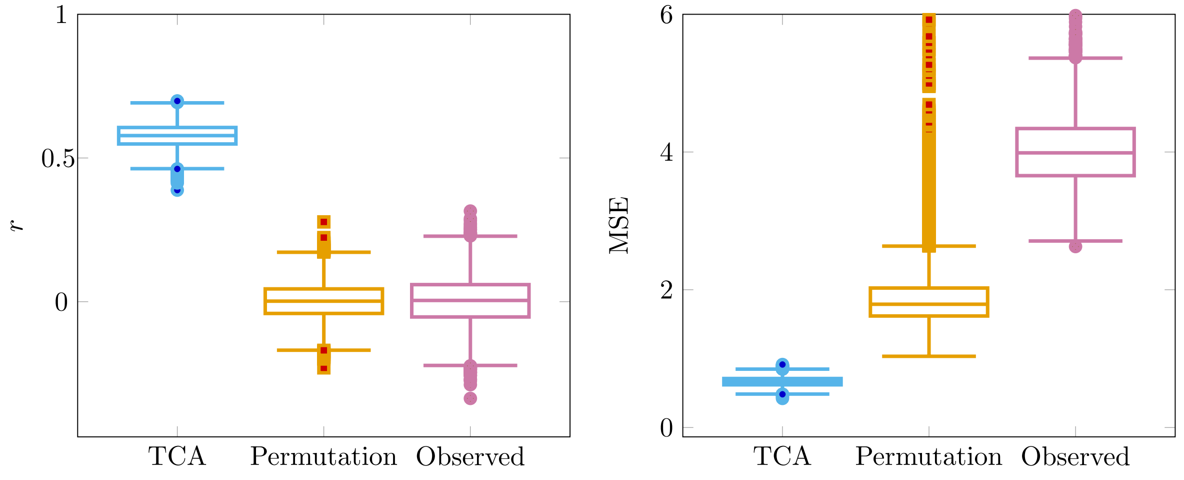

Figure 15: Reconstructing three-dimensional observation- and source-specific values from two-dimensional input across ten simulated data sets ($n = 250, m = 250, k = 3, \tau = 0.01$). Three approaches were evaluated in capturing the observation-specific values for each feature $j$ and source $h$ (i.e. $z_{hj}$): TCA, TCA after permuting the observed two-dimensional data matrix ("Permutation") and directly using the observed data matrix ("Observed"). For each of the evaluated approaches, we present the distribution of the linear correlation between $z_{hj}$ and its estimate $\hat{z}_{hj}$ across all $h, j$ (in the left) and the distribution of the MSE between $z_{hj}$ and its estimate $\hat{z}_{hj}$ across all $h, j$ (in the right).

permuting $X$ (independent permutation of each row of the matrix). In addition, for each vector $z_{hj}$, we also measured to what extent its information can be captured by $x_j = (x_{1j}, ..., x_{nj})^T$, the observed levels in the $j$-th feature of $X$. We observed that TCA could effectively reconstruct a substantial portion of the information in the $\{z_{hj}\}$ vectors, far outperforming the baseline measurements (Figure 15). We further verified the robustness of TCA by varying the parameters of the simulation across a wide range of values (Figure 16).

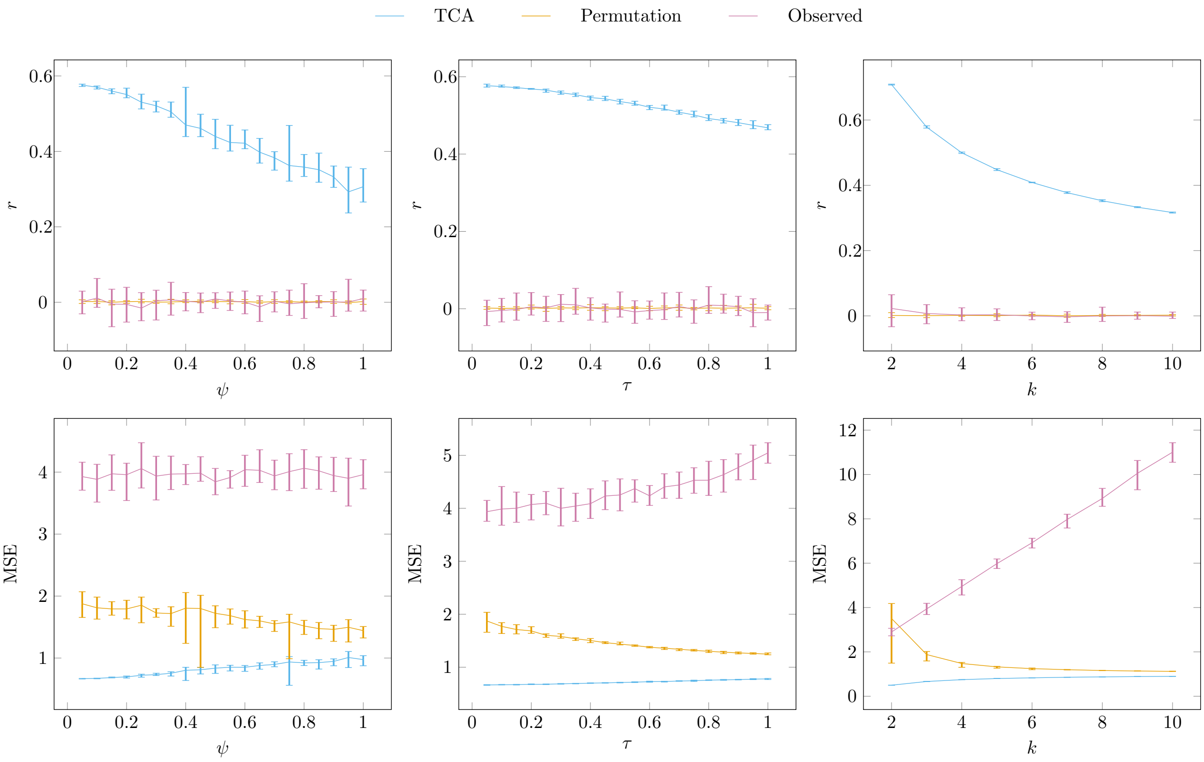

Figure 16: Reconstructing three-dimensional observation- and source-specific values from two-dimensional input in simulated data ($n = 250, m = 250$) while varying the parameters of the simulation. Data was simulated under three scenarios: increasing level of i.i.d. noise added to $W$ ($\psi$), increasing level of the i.i.d. component of variation added on top of $X$ ($\tau$) and increasing number of sources in the data ($k$). Three approaches were evaluated in capturing the observation-specific values for each feature $j$ and source $h$ ($z_{hj}$): TCA, TCA after permuting the observed data ("Permutation") and directly using the observed data ("Observed"). For each of the approaches and for each of the evaluated parameters, we present the median linear correlation between $z_{hj}$ and its estimate $\hat{z}_{hj}$ across all $h, j$ and across ten simulated data sets (top panel) and the median MSE between $z_{hj}$ and its estimate $\hat{z}_{hj}$ across all $h, j$ and across ten simulated data sets (bottom panel).

##### 2.6 Inferring the parameters of the model

In order to estimate the $\{z_{hj}^i\}$ values, the TCA algorithm requires knowledge of the parameters in (1) to (4). Since $X_{ij}$ is essentially a function of $Z_{1j}^i, ..., Z_{kj}^i$, we can use its assumed distribution for estimating all of the parameters in the model. More specifically, following the model in (3)-(4), we note that:

$$X_{ij} \sim N\left((c_i^{(2)})^T \delta_j + \sum_{h=1}^{k} w_{hi}\left(\mu_{hj} + (c_i^{(1)})^T \gamma_h^j\right), \sum_{h=1}^{k} w_{hi}^2 \sigma_{hj}^2 + \tau^2\right) \tag{17}$$

We can therefore take an ML approach for estimating the parameters of the model from the observed data matrix $X$. In practice, we require an initial estimate of $W$ as an input for the optimization. Such an estimate can be obtained by either using a reference-based approach [4] or a reference-free semi-supervised approach [8]. Given an estimate of $W$, we can then estimate the rest of the parameters in the model, and given estimates for the rest of the parameters in the model, we can update the estimate of $W$. We perform this alternating optimization procedure until convergence. Since we assume that different individuals are independent, updating $W$ requires us to solve a set of $n$ relatively easy optimization problems, each with $k$ parameters, while satisfying the constraints in (5) and (6); we solve this numerically using a standard non-linear optimization procedure. Below, we describe the optimization of the rest of the parameters of the model given $W$ (or an estimate of $W$).

Given $W$ and the variances $\tau, \sigma_j = (\sigma_{1j}, ..., \sigma_{kj})^T$, ML solution for $\mu_j = (\mu_{1j}, ..., \mu_{kj})^T$, $\delta_j$, $\{\gamma_h^j\}_{h=1}^k$ for feature $j$ is given by solving the following constrained regression problem:

$$\hat{\mu}_j, \hat{\delta}_j, \{\hat{\gamma}_h^j\}_{h=1}^k = \underset{\mu_j, \delta_j, \{\gamma_h^j\}_{h=1}^k}{\operatorname{argmin}} \sum_{i=1}^{n}\left(\tilde{x}_{ij} - \sum_{h=1}^{k} \tilde{w}_{hi}\mu_{hj} - \sum_{l=1}^{p_2} \tilde{c}_{li}^{(2)} \delta_{jl} - \sum_{l=1}^{p_1}\sum_{h=1}^{k} \tilde{c}_{lih}^{(1)} \gamma_h^j\right)^2 \tag{18}$$

$$\text{s.t.} \quad \forall 1 \leq j \leq m : \mu_{hj} \in [0,1] \tag{19}$$

where

$$\tilde{x}_{ij} = \frac{x_{ij}}{\sqrt{\sum_{l=1}^{k} w_{li}^2 \sigma_{lj}^2 + \tau^2}} \quad (20)$$

$$\tilde{w}_{ih} = \frac{w_{hi}}{\sqrt{\sum_{l=1}^{k} w_{li}^2 \sigma_{lj}^2 + \tau^2}} \quad (21)$$

$$\tilde{c}_{lih}^{(1)} = \frac{w_{ih} c_{li}^{(1)}}{\sqrt{\sum_{d=1}^{k} w_{di}^2 \sigma_{dj}^2 + \tau^2}} \quad (22)$$

$$\tilde{c}_{li}^{(2)} = \frac{c_{li}^{(2)}}{\sqrt{\sum_{h=1}^{k} w_{hi}^2 \sigma_{hj}^2 + \tau^2}} \quad (23)$$

and where $\delta_{jl}$ is the $l$-th entry of the vector $\delta_j$, and $c_{li}^{(1)}, c_{li}^{(2)}$ are the $l$-th covariate of individual $i$ in $C^{(1)}$ and in $C^{(2)}$, respectively. The constraints in (19) reflect the fact that methylation levels are bounded to the range $[0, 1]$, which means the mean levels should be also bounded to that range. We note that in principle we should also constrain the effects contributed by $\delta^j, \{\gamma_h^j\}_{h=1}^k$, in order to make sure that the total estimated methylation levels do not fall out of the range $[0, 1]$. In practice, in real data, these additional constraints may result with less accurate estimates. This problem can be solved efficiently using quadratic programming.

Since $\tau, \sigma_j$ are typically unknown, we perform an alternative optimization procedure as follows. We start by finding initial estimates for $\delta^j, \{\gamma_h^j\}_{h=1}^k$, by assuming that $\sigma_{1j} = ... = \sigma_{kj}, \tau = 0$. Under these conditions, the solution to the optimization problem in (18) is now independent of $\sigma_j, \tau$. Specifically, for obtaining an initial estimate of $\mu_j, \delta^j, \{\gamma_h^j\}_{h=1}^k$, we solve the problem in (18) while setting

$$\tilde{x}_{ij} = \frac{x_{ij}}{\|w_i\|_2} \quad (24)$$

$$\tilde{w}_{hi} = \frac{w_{hi}}{\|w_i\|_2} \quad (25)$$

$$\tilde{c}_{lih}^{(1)} = \frac{w_{hi} c_{lih}^{(1)}}{\|w_i\|_2} \quad (26)$$

$$\tilde{c}_{li}^{(2)} = \frac{c_{li}^{(2)}}{\|w_i\|_2} \quad (27)$$

Once we obtain $\hat{\mu}_j, \hat{\delta}_j, \{\hat{\gamma}_h^j\}_{h=1}^k$, we can fix them and estimate $\sigma_j, \tau$ using any hill climbing algorithm (and then repeat until convergence). In practice, for learning $\sigma_j, \tau$ we perform another alternating optimization procedure as follows. We first assume $\tau$ to be unique for each site and estimate for each site $j$ separately initial estimates of $\sigma_j, \tau$. Then, we re-estimate $\tau$ using the entire data and the estimates of $\{\sigma_j\}$ from all sites, and finally, we re-estimate $\sigma_j$ for each site $j$ using the updated estimate of $\tau$.

Notably, the number of parameters we need to estimate in our model is very large compared with the number of data points available for inference. However, for each set of constant number of parameters that we estimate, we use $n$ data points. For instance, for estimating the parameters $\mu_j, \delta_j, \{\gamma_h^j\}_{h=1}^k$ for site $j$ (a constant set of $k(p_1 + 1) + p_2$ parameters), we use $n$ data points.

#### 2.7 Testing a phenotype for cell-type-specific associations

TCA allows us to estimate cell-type-specific methylation levels for each individual in the data. In principle, such estimates can then be used for running a cell-type-specific EWAS by testing the estimates of a particular cell type for association with a phenotype of interest (or for running a joint test for several cell types by using their estimated cell-type-specific methylation levels jointly). However, for the application of association testing, we suggest an alternative one-step approach instead of the more straightforward two-steps approach.

We model the phenotype of interest as potentially affected by cell-type-specific methylation levels, and use the conditional distribution of the phenotype given the observed data in $X$. Effectively, this allows us to integrate over all the potential values of the $\{z_{hj}^i\}$ individual and cell-type-specific levels. In addition to taking into account covariates that may affect the methylation levels, as described in (1) and in (3), we also consider potential direct effects of other (or the same) covariates on the phenotype.

#### 2.8 Joint test for effect sizes in all cell types

Let $Y \in \mathbb{R}^{n \times 1}$ be a quantitative phenotype of interest, where $Y_i$ corresponds to the phenotypic level of sample $i$, and let $C^{(3)} \in \mathbb{R}^{p_3 \times n}$ be a matrix of $p_3$ covariates potentially affecting the phenotype (may also include an intercept term), we assume the following model:

$$Y_i = (c_i^{(3)})^T \alpha + \sum_{h=1}^{k} \beta_{hj} Z_{hj}^i + e_i \tag{28}$$

$$e_i \sim N(0, \phi^2) \tag{29}$$

where $\beta_{1j}, ..., \beta_{kj}$ are the effect sizes of the $k$ different cell types in site $j$. Recall the model of $X_{ij}$ in (3)-(4), using (12) we get

$$Y_i | X_{ij} \sim N\left(\tilde{\mu}_1 + \frac{COV(X_{ij}, Y_i)}{\tilde{\sigma}_2^2}(x_{ij} - \tilde{\mu}_2), \tilde{\sigma}_1^2 - \frac{COV(X_{ij}, Y_i)^2}{\tilde{\sigma}_2^2}\right) \tag{30}$$

where

$$\tilde{\mu}_1 = E(Y_i), \tilde{\sigma}_1^2 = V(Y_i) \tag{31}$$

$$\tilde{\mu}_2 = E(X_{ij}), \tilde{\sigma}_2^2 = V(X_{ij}) \tag{32}$$

Different individuals are assumed to be independent (both in their phenotypic and their cell-type-specific methylation levels), and $COV(y_i, X_{tj}) = 0$ for any $t \neq i$.

Note that

$$\begin{aligned}
COV(X_{ij}, Y_i) &= E(Y_i X_{ij}) - E(Y_i)E(X_{ij}) \\
&= E\left(\left((c_i^{(3)})^T \alpha + \sum_{h=1}^{k} \beta_{hj} Z_{hj}^i + e\right)\left((c_i^{(2)})^T \delta^j + \sum_{h=1}^{k} w_{hi} Z_{hj}^i + \epsilon\right)\right) \\
&- \left((c_i^{(3)})^T \alpha + \sum_{h=1}^{k} \beta_{hj} E(Z_{hj}^i)\right)\left(c_i^{(2)} \delta^j + \sum_{h=1}^{k} w_{hi} E(Z_{hj}^i)\right) \\
&= (c_i^{(3)})^T \alpha \sum_{h=1}^{k} w_{hi} E(Z_{hj}^i) + (c_i^{(2)})^T \delta_j \sum_{h=1}^{k} \beta_{hj} E(Z_{hj}^i) + E\left(\sum_{h=1}^{k} \beta_{hj} Z_{hj}^i \sum_{h=1}^{k} w_{hi} Z_{hj}^i\right) \\
&- (c_i^{(3)})^T \alpha \sum_{h=1}^{k} w_{hi} E(Z_{hj}^i) - (c_i^{(2)})^T \delta_j \sum_{h=1}^{k} \beta_{hj} E(Z_{hj}^i) - \sum_{h=1}^{k} \beta_{hj} E(Z_{hj}^i) \sum_{h=1}^{k} w_{hi} E(Z_{hj}^i) \\
&= \sum_{h=1}^{k} w_{hi} \beta_{hj} E((Z_{hj}^i)^2) - \sum_{h=1}^{k} w_{hi} \beta_{hj} E(Z_{hj}^i)^2 \\
&= \sum_{h=1}^{k} w_{hi} \beta_{hj} \sigma_{hj}^2
\end{aligned} \tag{33}$$

Therefore, we get

$$Y_i | X_{ij} = x_{ij} \sim N\left(\tilde{\mu}_{ij}, \tilde{\sigma}_{ij}^2\right) \tag{34}$$

where

$$\tilde{\mu}_{ij} = (c_i^{(3)})^T\alpha + \sum_{h=1}^{k}\beta_{hj}\left(\mu_{hj} + (c_i^{(1)})^T\gamma_h^j + \frac{w_{hi}\sigma_{hj}^2\tilde{x}_{ij}}{\tau^2 + \sum_{l=1}^{k} w_{li}^2\sigma_{lj}^2}\right) \quad (35)$$

$$\tilde{x}_{ij} = x_{ij} - (c_i^{(2)})^T\delta_j - \sum_{l=1}^{k} w_{li}(\mu_{lj} + (c_i^{(1)})^T\gamma_l^j) \quad (36)$$

$$\tilde{\sigma}_{ij}^2 = \phi^2 + \sum_{h=1}^{k}\beta_{hj}^2\sigma_{hj}^2 - \frac{\left(\sum_{h=1}^{k}\beta_{hj}w_{hi}\sigma_{hj}^2\right)^2}{\tau^2 + \sum_{h=1}^{k} w_{hi}^2\sigma_{hj}^2} \quad (37)$$

Using the distributions $Y_i|X_{ij} = x_{ij}$ for each individual $i$, we can now consider the following hypothesis testing for site $j$:

$$H_0 : \beta_{1j} = ... = \beta_{kj} = 0 \quad (38)$$

$$H_1 : \exists h.\beta_{hj} \neq 0 \quad (39)$$

This formulation essentially tests the particular site under test $j$ for association with the phenotype by considering the joint contribution of all cell-type-specific effects. Alternatively, we can look for cell-type-specific effects of a subset of the cell types.

##### 2.9 Marginal test for the effect size of a particular cell type

Consider the following model:

$$Y_i = (c_i^{(3)})^T\alpha + \beta_{hj}Z_{hj}^i + e_i \quad (40)$$

$$e_i \sim N(0, \phi^2) \quad (41)$$

where $\beta_{hj}$ is the effect size of a particular cell type $h$. Similarly as before, we get:

$$Y_i|X_{ij} = x_{ij} \sim N\left(\tilde{\mu}_{ij}, \tilde{\sigma}_{ij}^2\right) \quad (42)$$

where

$$\tilde{\mu}_{ij} = (c_i^{(3)})^T\alpha + \beta_{hj}\left(\mu_{hj} + (c_i^{(1)})^T\gamma_h^j + \frac{w_{hi}\sigma_{hj}^2\tilde{x}_{ij}}{\tau^2 + \sum_{l=1}^{k} w_{li}^2\sigma_{lj}^2}\right) \quad (43)$$

$$\tilde{x}_{ij} = x_{ij} - (c_i^{(2)})^T\delta_j - \sum_{l=1}^{k} w_{li}(\mu_{lj} + (c_i^{(1)})^T\gamma_l^j) \quad (44)$$

$$\tilde{\sigma}_{ij}^2 = \phi^2 + \beta_{hj}^2\left(\sigma_{hj}^2 - \frac{w_{hi}^2\sigma_{hj}^4}{\tau^2 + \sum_{l=1}^{k} w_{li}^2\sigma_{lj}^2}\right) \quad (45)$$

Using the distributions $Y_i|X_{ij} = x_{ij}$ for each individual $i$, we can now consider the following hypothesis testing for site $j$:

$$H_0 : \beta_{hj} = 0 \tag{46}$$

$$H_1 : \beta_{hj} \neq 0 \tag{47}$$

We calculate p-values for both the joint test and the marginal test using a generalized likelihood-ratio test. The null model can be fitted using standard ML estimators. For the alternative model, given the estimates for a particular site $j$, $\Theta_j = (\mu_j, \sigma_j, W, \tau, \Gamma_j, \delta_j)$, and given the observed data $Y, X_j, C^{(1)}, C^{(2)}, C^{(3)}$, the parameters $\alpha = (\alpha_1, ..., \alpha_p), \phi$ and $\beta_j = (\beta_{1j}, ..., \beta_{kj})$ (in a marginal test for cell type $h$ only the estimate of $\beta_{hj}$ is needed) can be estimated using ML. In practice, we do that by numerically maximizing the log likelihood of the conditional distribution using a standard non-linear optimization procedure.

Throughout our experiments in the paper, we observed that TCA, albeit powerful, resulted in a deflation in the test statistic under the null, leading it to be an over-conservative test. This behavior may be explained by the optimization procedure we apply. Specifically, an appropriate application of the generalized-likelihood ratio test we use relies upon using ML estimates of the parameters in the TCA model. In our case, we achieve ML estimates under the null model, however, in general, we do not achieve ML estimates under the alternative model for two reasons. First, our optimization procedure involves a non-convex optimization, which is not guaranteed to yield global optimum, and second, for computational convenience, we leverage only the bulk methylation data ($X$) in learning the parameters of the TCA model. The latter is not optimal since in principle the phenotypic data ($Y$) provides more information about the parameters of the model. As a result, the estimates under the alternative hypothesis are not ML estimates, which leads to a lower likelihood of the alternative model and therefore to a deflation in the test statistic of the generalized-likelihood ratio test (and thus the test is over-conservative).
